## Supplemental figures for "Neuronal Identity is Not Static—An Input-Driven Perspective"

**This PDF file includes:**

Figures S1 to S6

### 1 Waveform and Action Potential properties manifold comparison

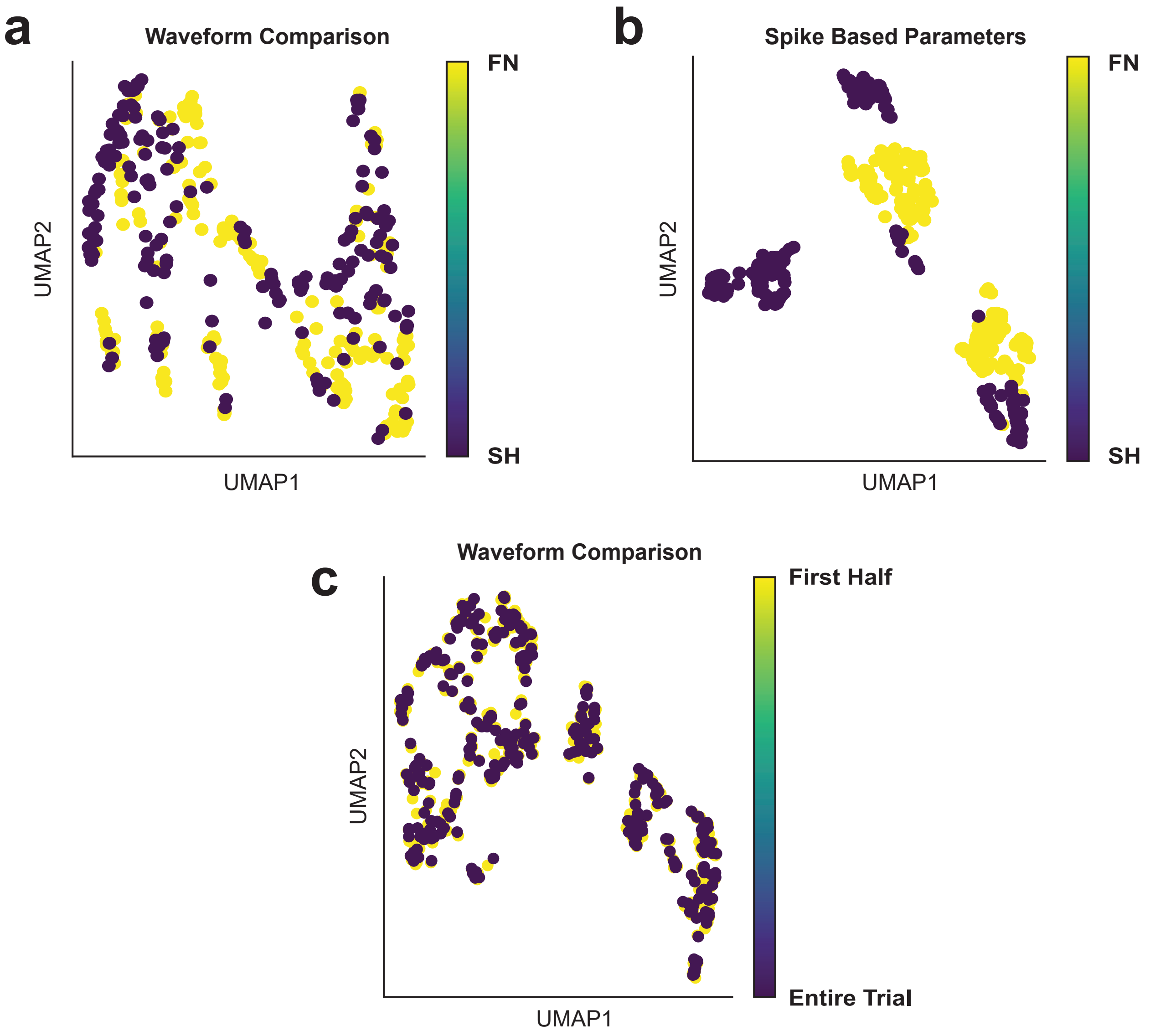


(**a**) Overlaid UMAP representation FN and SH waveforms from 186 neuron used in classification. The waveform shapes are different between SH and FN protocols. (**b**) FN and SH Action potential parameters used for classification projected together on the same space. The Action potential properties are different between FN and SH protocols. The SH Action potential properties show a bigger spread than FN properties. (**c**) UMAP projection of averaged Waveform comparison between the first half and the entire trial. The average waveform shape doesn't change as a result of trial length.

#### 2 Louvain vs Ensemble Clustering for Graphs (ECG) algorithm comparison

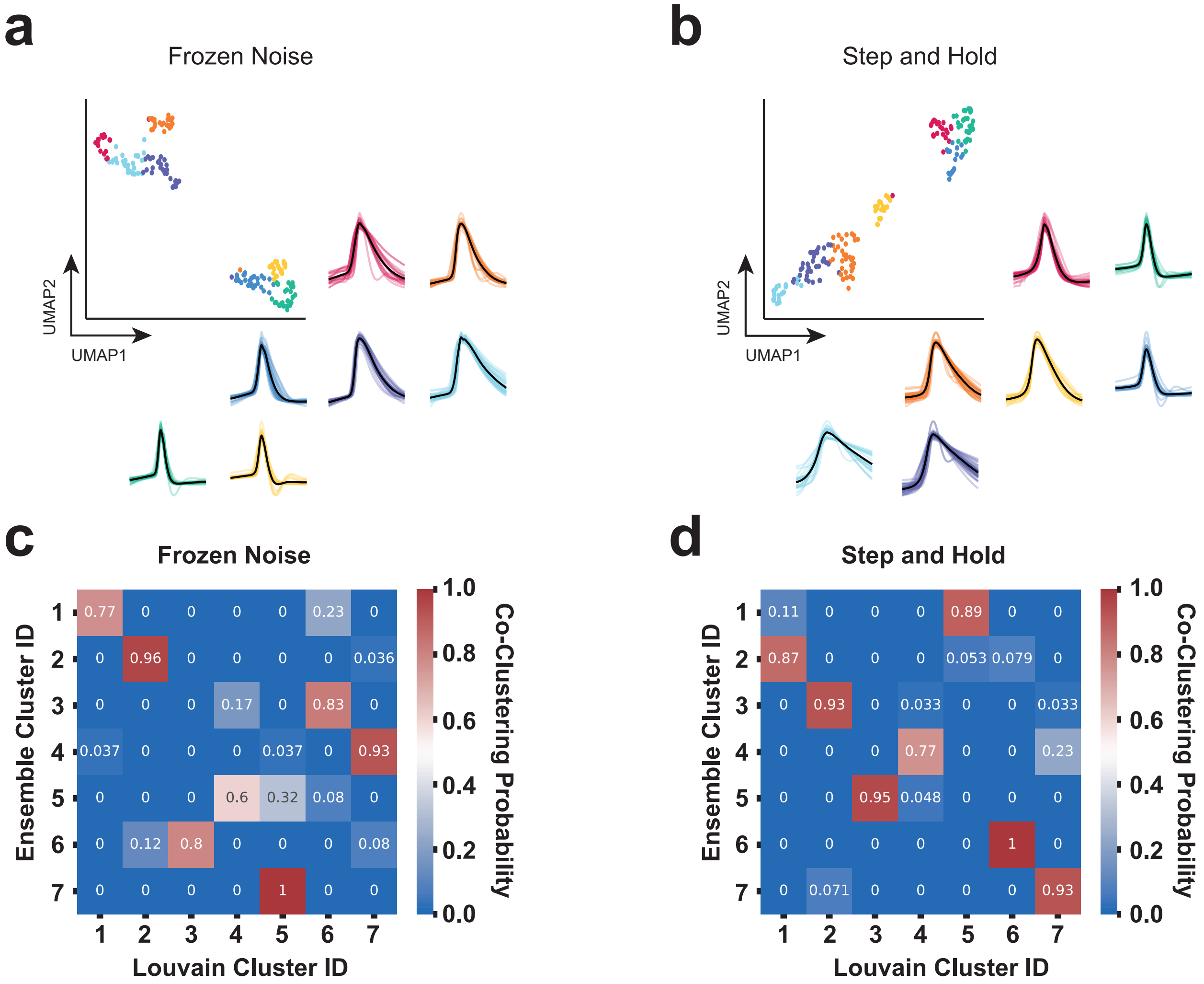


(**a**) UMAP embedding of FN waveforms colored with cluster labels found using Ensemble clustering, with original waveforms in the same color as the respective color. 7 clusters were observed, the same number as in the case of the Louvain community method. (**b**) UMAP embedding of SH waveforms colored with cluster labels found using Ensemble clustering, with original waveforms in the same color as the respective color. 7 clusters were observed, the same number as in the case of the Louvain community method. (**c**) Heatmap showing the correspondence between Louvain and Ensemble clustering for graph on FN waveforms. Clusters using the Louvain community detection algorithm show a high correspondence with the clusters obtained using Ensemble clustering for the graph method. (**d**) Heatmap showing the correspondence between Louvain and Ensemble clustering for graph on SH waveforms. Clusters using the Louvain community detection algorithm show a high correspondence with the clusters obtained using Ensemble clustering for the graph method.

### 3 Cluster stability after leaving one attribute out at a time

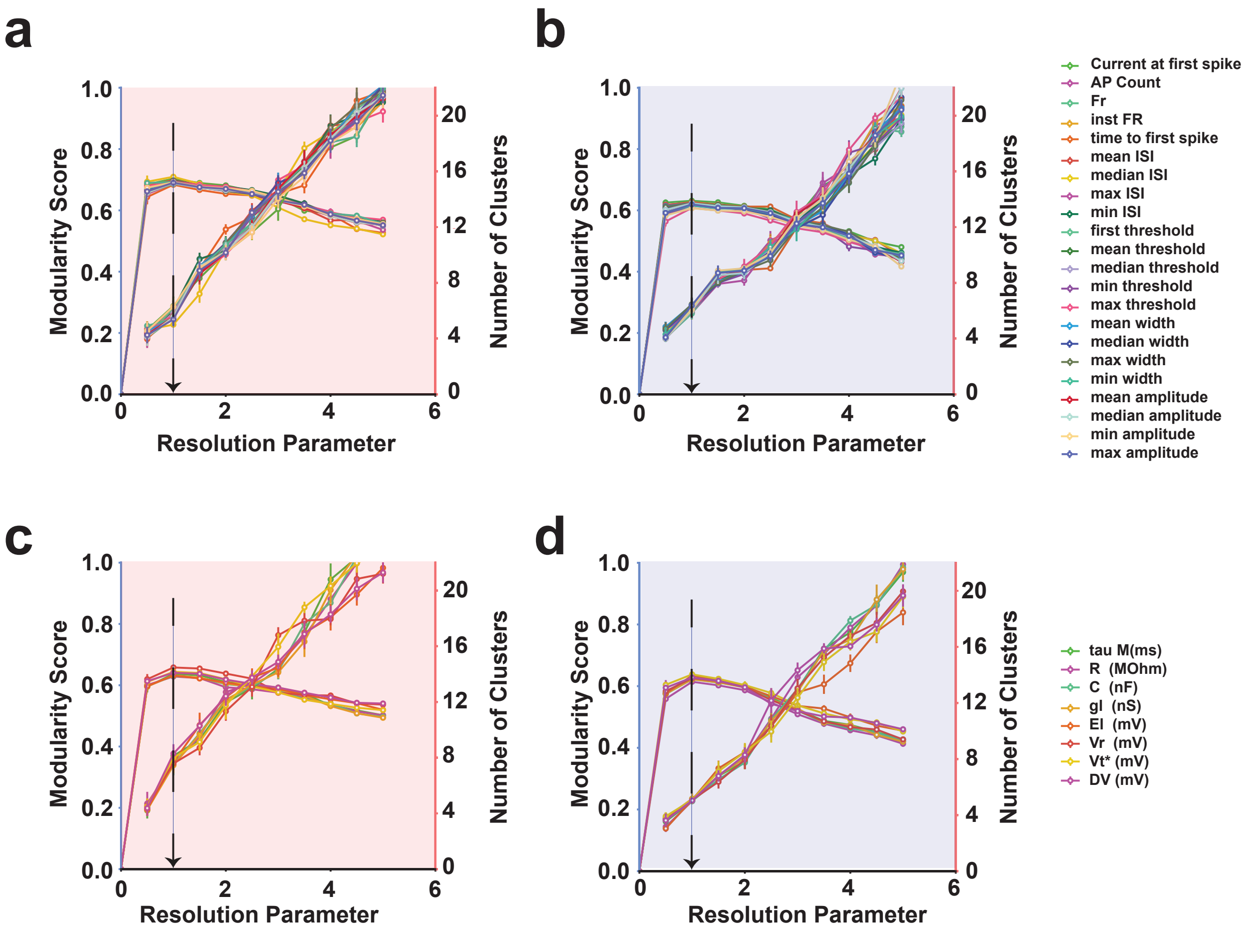


(**a-b**) Stability of clusters for action potential attributes leaving one attribute out for excitatory and Inhibitory sets. The clustering was performed 25 times with random 80% samples for resolution attributes ranging from 0.0 to 1, the mean and standard deviation of the resulting number of clusters and modularity score are plotted. For the chosen resolution parameter (1.0), the cluster numbers fluctuate between 5-7. The cluster number fluctuates more for the excitatory set (left) than the Inhibitory set (right) (**c-d**) Stability of clusters for biophysical attributes leaving one attribute out for excitatory and Inhibitory sets. The clustering was performed 25 times with random 90% samples for resolution parameters ranging from 0.0 to 1, the mean and standard deviation of the resulting number of clusters and modularity score are plotted. For the chosen resolution parameter (1.0), the excitatory population fluctuates between 6-8. On the contrary, the Inhibitory population doesn't show any fluctuation.

### 4 Diversity of firing rate and AP half-width

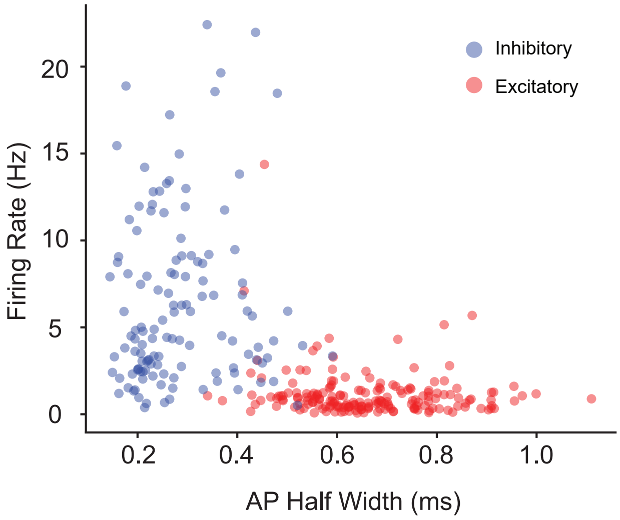


Firing rate vs AP half-width between excitatory (red) and inhibitory (blue) populations. The excitatory population has a lower firing rate and higher AP width. The Inhibitory population has a lower AP width and higher firing rate.

### 5 STA Heterogeneity

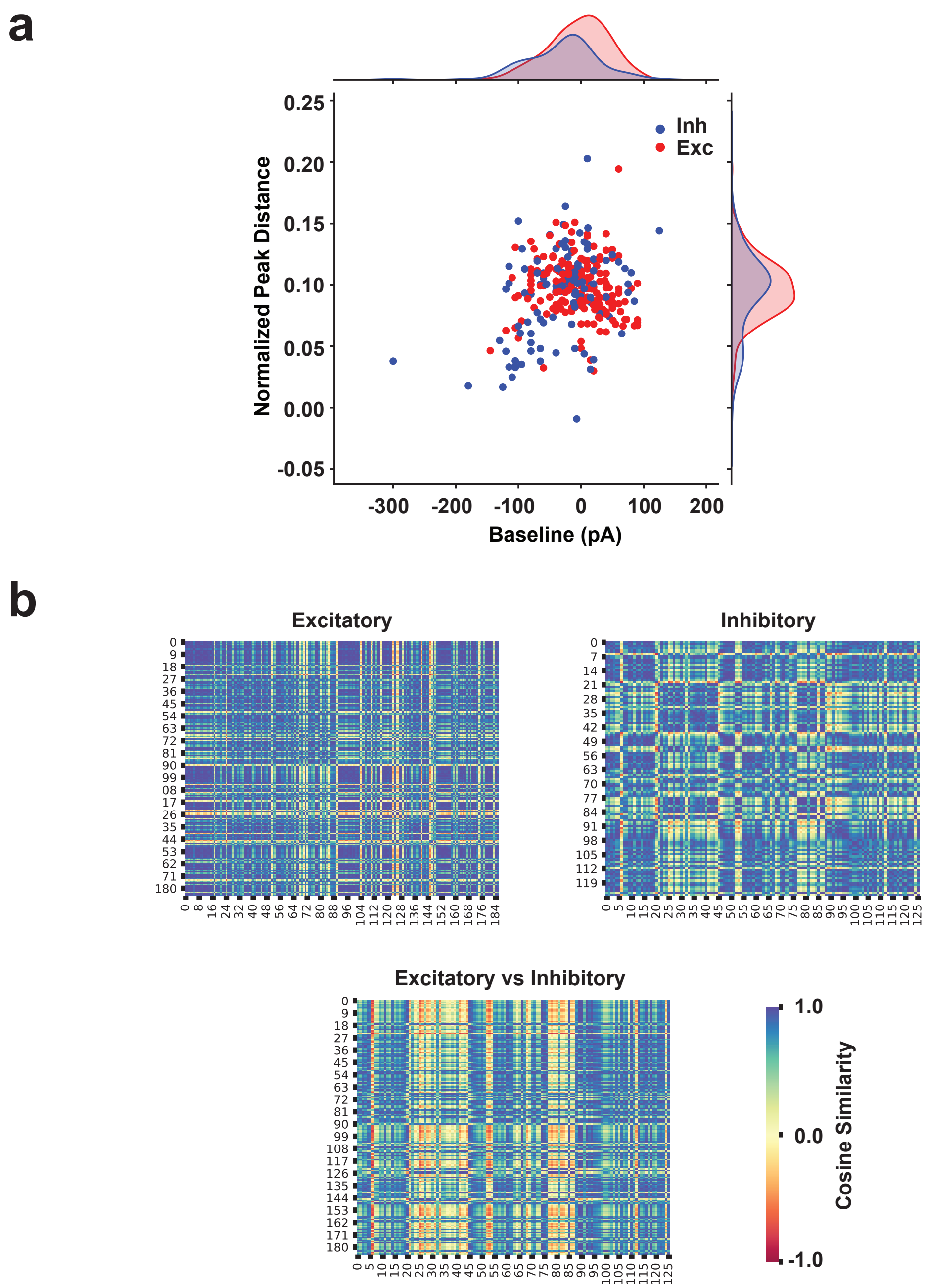


(**a**) The scatter plot shows the baseline added to the theoretical input vs the Peak distance of the STA. The excitatory population has a higher baseline and higher peak distance. The variance for both baseline and peak distance seems higher for the excitatory population than inhibitory population. (**b**) The STA cosine similarity is for the excitatory population (top left), the STA cosine similarity is for the inhibitory population (top right), and the STA cosine similarity is between the excitatory and inhibitory populations. The excitatory population has higher similarity than the inhibitory population.

### 6 Cluster assignment likelihood across attributes

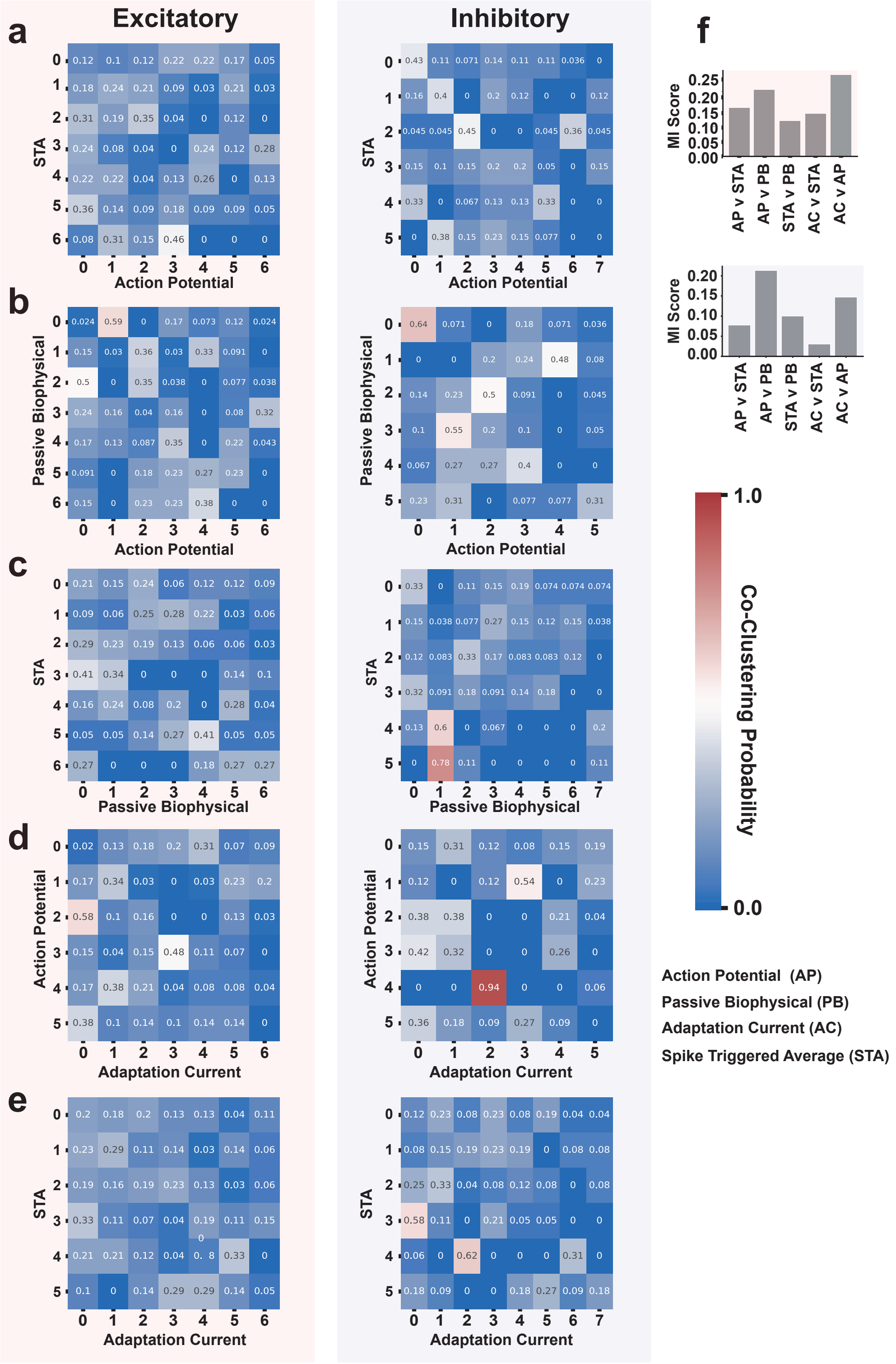


(**a-e**) Heatmap showing the likelihood for neuron clustering in attribute 1 clustering in one of the clusters in attribute 2 for excitatory (left) and inhibitory (right) population. For each attribute

(Action potential, Passive biophysical, STA, Adaptation current ($\eta$), none of the clusters show a high likelihood to be clustering in another attribute. (f) Cluster label comparison between modularity pairs using Mutual information score for excitatory and inhibitory population. None of the pairs reach the modularity of 0.25 showing that neurons cluster differently across attribute sets.
